## Supplementary Material for "Eukaryotic CRFK cells motion characterized with atomic force microscopy"

[b] Centro de Investigación en Sanidad Animal, INIA-CISA, Valdeolmos, Madrid, 28130, Spain.

Supplementary Fig. 1

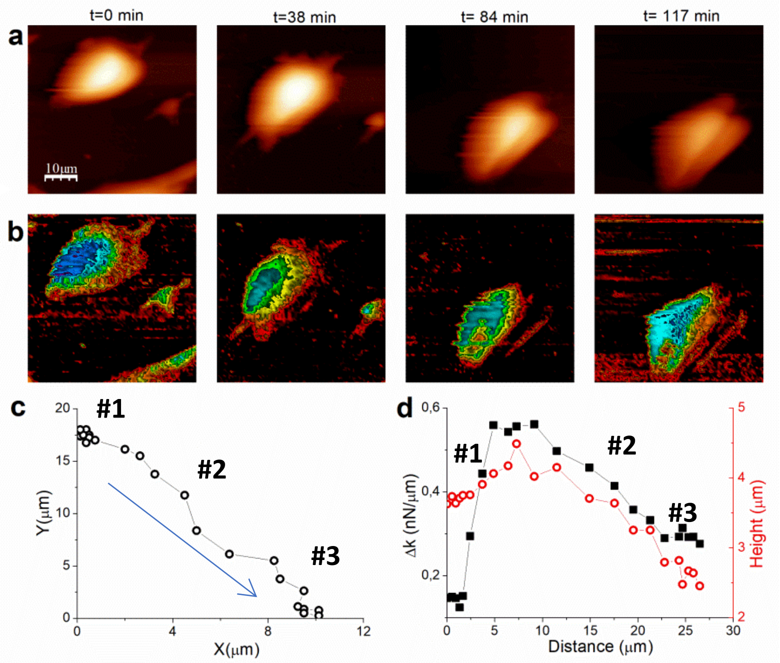


**Suppl. Fig. 1.** Series of **a)** topographic (Z-scale = 4 µm) and **b)** Spring constant k (Z-scale: 2 nN/µm), images (50 µm × 50 µm) recorded in real time for a single CRFK cell during its motion on glass surface for t = 0, 38, 84 and 117 min. respectively. **c**) Trajectory of the cell plotted as the position of the cell for the aligned images. **d)** Plot of $\Delta k_{rear\text{-}front}$ (black) and cell maximum height (red), as function of the cell's traveled distance.

Time-lapse AFM images were recorded in cells culture medium (DMEM) at 37 ºC. 22 successive images were collected at fixed time intervals of ≈ 6 minutes. We monitored the topography (**Suppl. Fig. 1a**) and spring constant (**Suppl. Fig. 1b**) response of the cell during its motion at glass slide surface. During the first 30 minutes, the cell is almost stopped (**Suppl. Fig. 1c**). This phase 1 (#1) is characterized by constant Δk≈ 0.1 nN/µm and height ≈ 3.5 µm (**Suppl. Fig. 1d**). The cells start moving when Δk increase up to ≈ 0.55 nN/µm and cell height reaches ≈ 5 µm. During this linear motion (#2), the cell travels ≈ 10 µm, Δk and cell height decrease linearly to ≈ 0.3 nN/µm and ≈ 3.5 µm respectively until the cell stops again in a phase 3 (#3) where Δk and the cell height keep unchanged again.

Supplementary Fig. 2

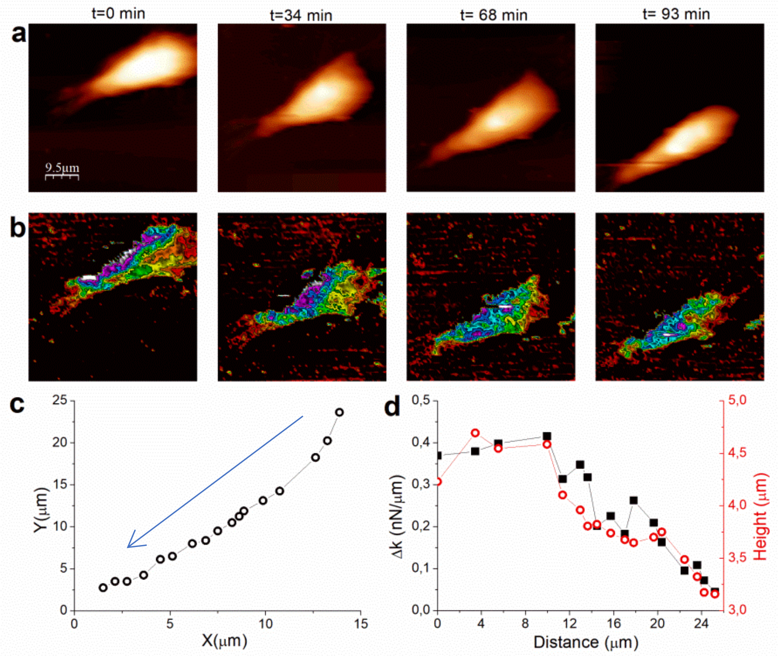


**Suppl. Fig. 2.** Series of **a)** topographic (Z-scale = 4 µm) and **b)** Spring constant k (Z-scale: 3.6 nN/µm), images (50 µm × 50 µm) recorded in real time for a single CRFK cell during its motion on glass surface for t = 0, 28, 57 and 86 min. respectively. **c**) Trajectory of the cell plotted as the position of the cell for the aligned images. **d)** Plot of $\Delta k_{rear\text{-}front}$ (black) and cell maximum height (red), as function of the cell's traveled distance.

Time-lapse AFM images were recorded in cells culture medium (DMEM) at 37 ºC. 17 Successive images were collected at fixed time intervals of ≈ 6 minutes. We monitored the topography (**Suppl. Fig. 2a)** and spring constant **(Suppl. Fig. 2b)** response of the cell during its motion at glass slide surface. The cell travels 25 µm mainly from the upper right to the center of the image (**suppl. Fig. 2c, blue arrow**). During the linear motion, Δk and cell height decrease linearly from ≈ 0.4 to ≈ 0.05 nN/µm and from≈ 4.5 to 3.2 µm respectively (**suppl. Fig. 2d**). Surprisingly the cell does not move along its large axis and it rotates slightly counterclockwise while moving

Supplementary Fig. 3

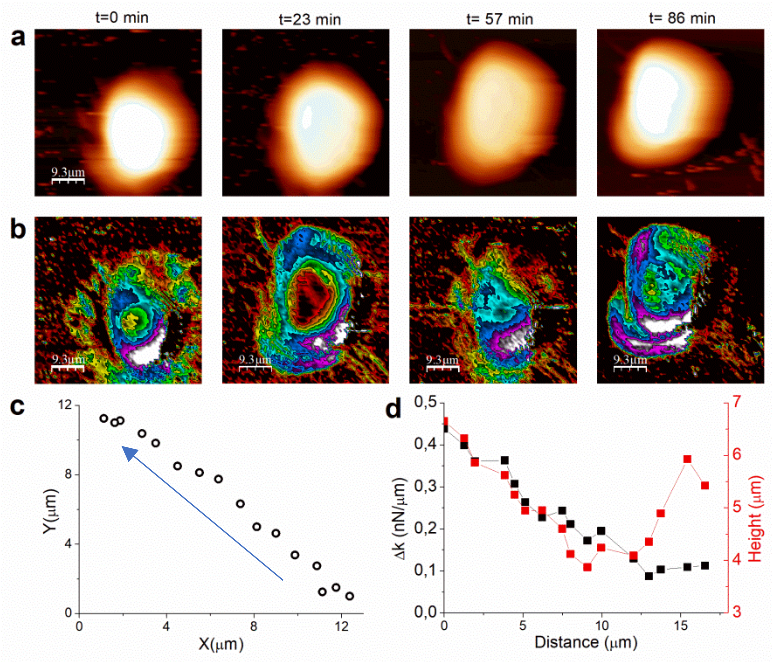


**Suppl. Fig. 3.** Series of **a)** topographic (Z-scale = 5 µm) and **b)** Spring constant k (Z-scale: 1.2 nN/µm), images (50 µm × 50 µm) recorded in real time for a single CRFK cell during its motion on glass surface for t = 0, 23, 57 and 86 min. respectively. **c**) Trajectory of the cell plotted as the position of the cell for the aligned images. **d)** Plot of $\Delta k_{rear\text{-}front}$ (black) and cell maximum height (red), as function of the cell's traveled distance.

Time-lapse AFM images were recorded in cells culture medium (DMEM) at 37 ºC. 16 successive images were collected at fixed time intervals of ≈ 6 minutes. We monitored the topography (**Suppl. Fig. 3a)** and spring constant **(Suppl. Fig. 3b)** response of the cell during its motion at glass slide surface. The cell shows a linear change in position traveling 17 µm from the bottom left to the upper right of the image (**suppl. Fig. 3c, blue arrow**). During the linear motion, Δk and cell height decrease linearly from ≈ 0.45 to ≈ 0.1 nN/µm and from≈ 6.5 to 4 µm respectively (**suppl. Fig. 3d**) until the cell nearly stop. Then the cell height increases again to ≈ 6 µm during the last 20 minutes of the experiment.

### Supplementary Fig. 4

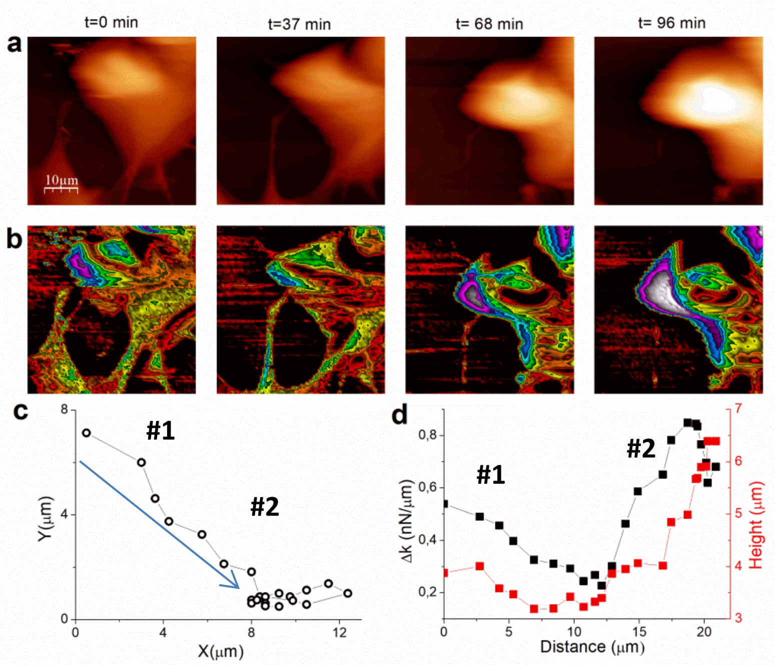


**Suppl. Fig. 4.** Series of **a)** topographic (Z-scale = 6 µm) and **b)** Spring constant k (Z-scale: 1.5 nN/µm), images (50 µm × 50 µm) recorded in real time for a single CRFK cell during its motion on glass surface for t = 0, 37, 68 and 96 min. respectively. **c**) Trajectory of the cell plotted as the position of the cell for the aligned images. **d)** Plot of $\Delta k_{rear\text{-}front}$ (black) and cell maximum height (red), as function of the cell's traveled distance.

Time-lapse AFM images were recorded in cells culture medium (DMEM) at 37 ºC. 22 successive images were collected at fixed time intervals of ≈ 6 minutes. We monitored the topography (**Suppl. Fig. 4a)** and spring constant **(Suppl. Fig. 4b)** response of the cell during its motion at glass slide surface. This cell is strongly connected to it neighboring cells from the upper left and the bottom of the image. During the first 50 minutes (blue arrow), the cell linearly travels 8 µm (**Suppl. Fig. 4c, #1**). Δk and cell height decrease linearly from ≈ 0.55 to 0.2 nN/µm and from≈ 4 to 3 µm respectively (**Suppl. Fig. 4d**). The second phase (#2) is more complex as the cell changes its direction. The cell is still moving but it also rotates counter clockwise to orientate following the X-axis then nearly stops. While rotating, Δk and cell height increases from ≈ 0.2 to 0.7 nN/µm and from≈ 3 to 6.5 µm respectively.

Supplementary Fig. 5

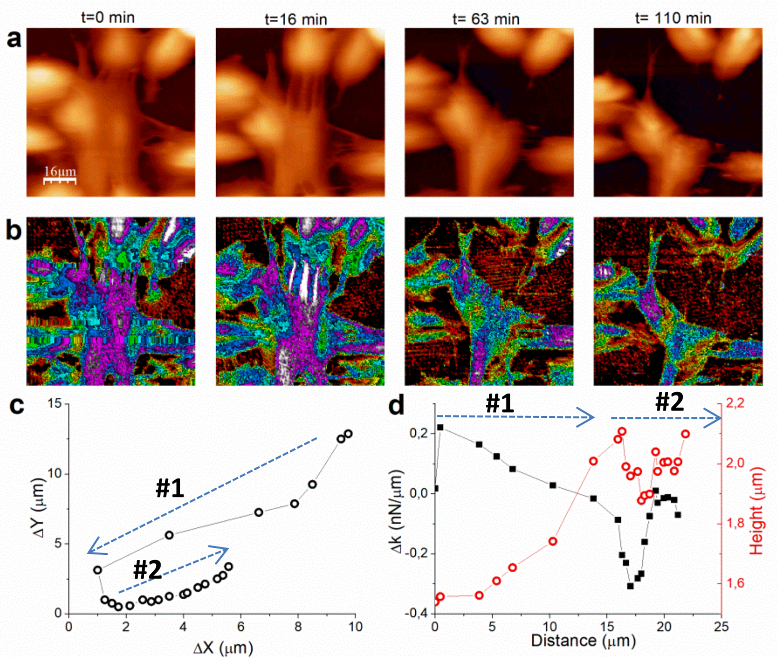


**Suppl. Fig. 5a)** Series of **a)** topographic (Z-scale = 4 µm) and **b)** Spring constant k (Z-scale: 2 nN/µm), images (80 µm ×80 µm) recorded in real time for a single CRFK cell during its motion on glass surface for t = 0, 16, 63 and 110 min. respectively. **c**) Trajectory of the cell plotted as the position of the cell for the aligned images. **d)** Plot of $\Delta k_{rear\text{-}front}$ (black) and cell maximum height (red), as function of the cell's traveled distance.

Time-lapse AFM images were recorded in cells culture medium (DMEM) at 37 ºC. 22 successive images were collected at fixed time intervals of ≈ 6 minutes. We monitored the topography (**Suppl. Fig. 5a)** and spring constant **(Suppl. Fig. 5b)** response of the cell during its motion at glass slide surface. We characterized the motion central cell (dotted blue circle), because it showed the largest changes in position during the measurements. The cell starts moving downward about 10 um and left (about 9 um), during 40 min (**Suppl. Fig.5.c, #1**). This movement is characterized by a decrease in the k cell junctions to the upper cell (**Suppl. Fig. 5b, Image 2, red arrow**). Then it stops and changes direction to move slower in the opposite direction during the rest of the experiment (# 2). During the phase 1 (#1), Δk (**Suppl. Fig. 5d, black curve)** is positive to ≈ 0.2 nN/µm and decreases linearly until the cell stops when Δk = 0 nN/µm. Cell height (**Suppl. Fig. 5d, red curve**) increases from ≈ 1.6 to 2.1 µm because the cell detaches from the upper cell when softer junctions break, leading to a change in cell spreading during motion. Then k reverses and becomes negative to Δk = -0.35 nN/µm during phase 2 (#2) as the cell changes its trajectory to move to the opposite direction. Cell height decreases when k dissymmetry increases emphasizing the link between cytoskeleton reorganization and cell morphology.

**Description of the additional Supplementary Movie.**

226 min JM-AFM of 40 images sequence of the CRFK cell motion after images correlation to correct for drift. Movie parameters: frame size: 73 µm. Full color Z-scale: 3 µm. Image acquisition speed: 10 Hz.
