## Supplementary figures and images for "Eukaryotic CRFK cells motion characterized with atomic force microscopy"

### Supplementary Figure 1

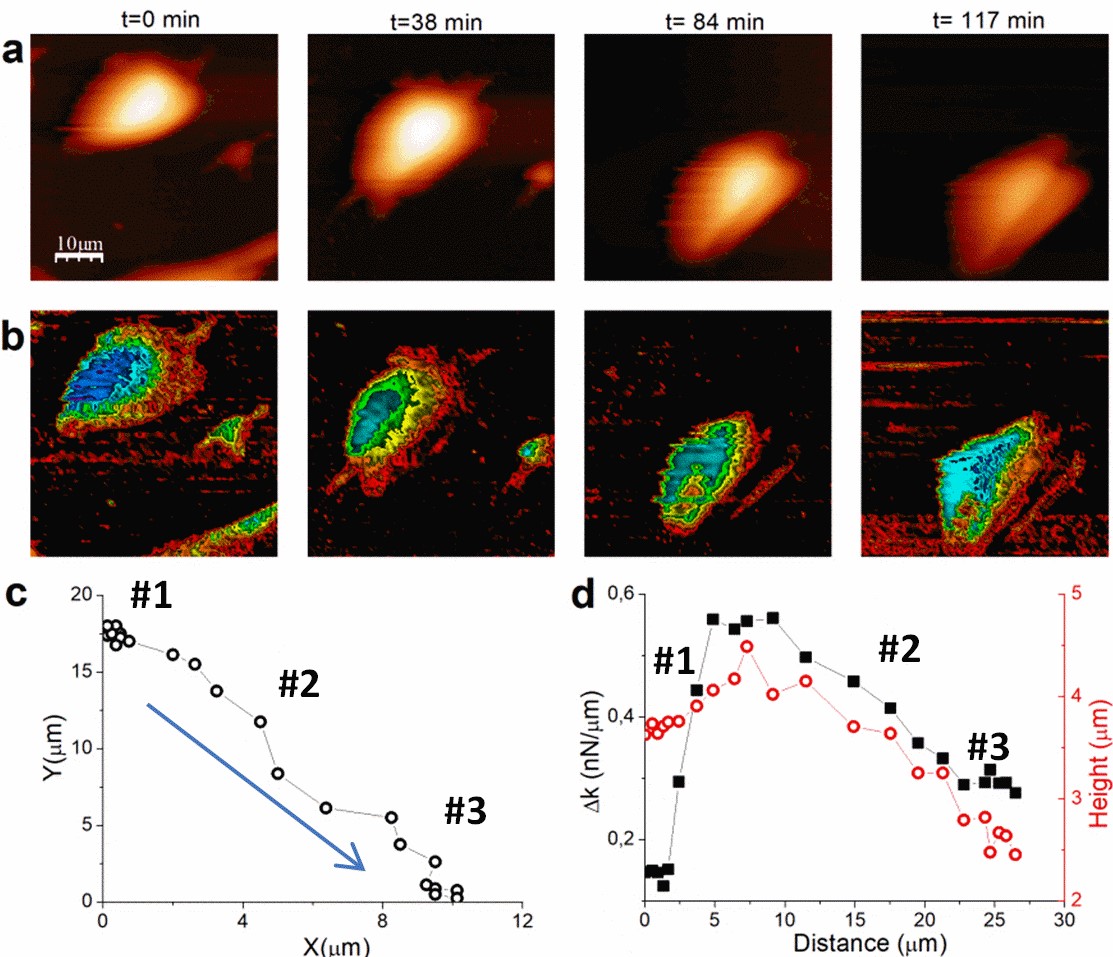

### Supplementary Figure 2

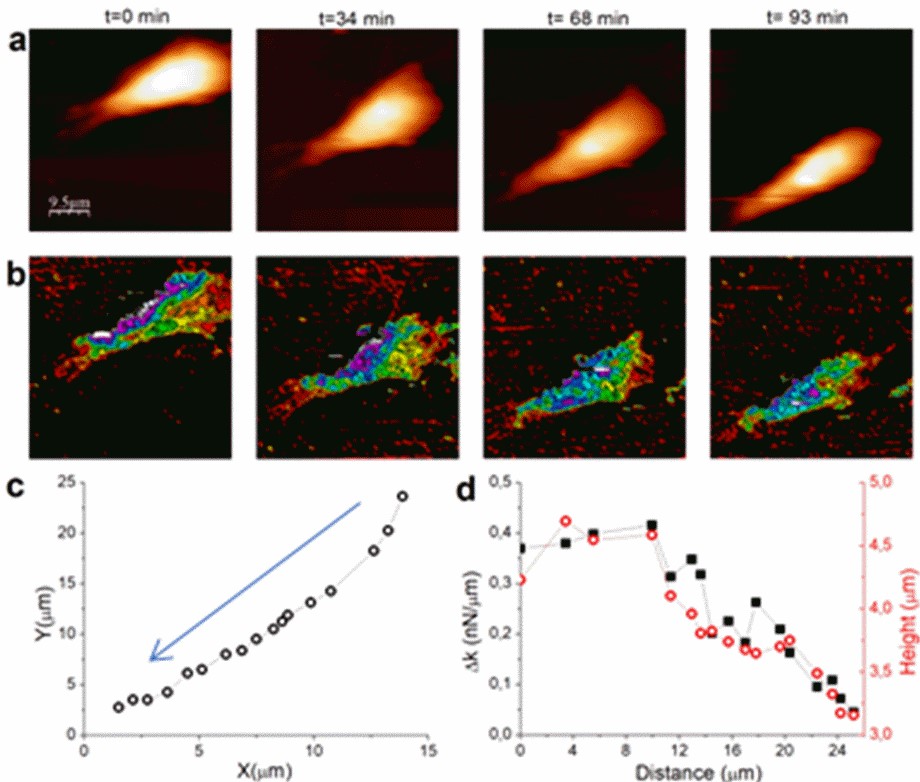

### Supplementary Figure 3

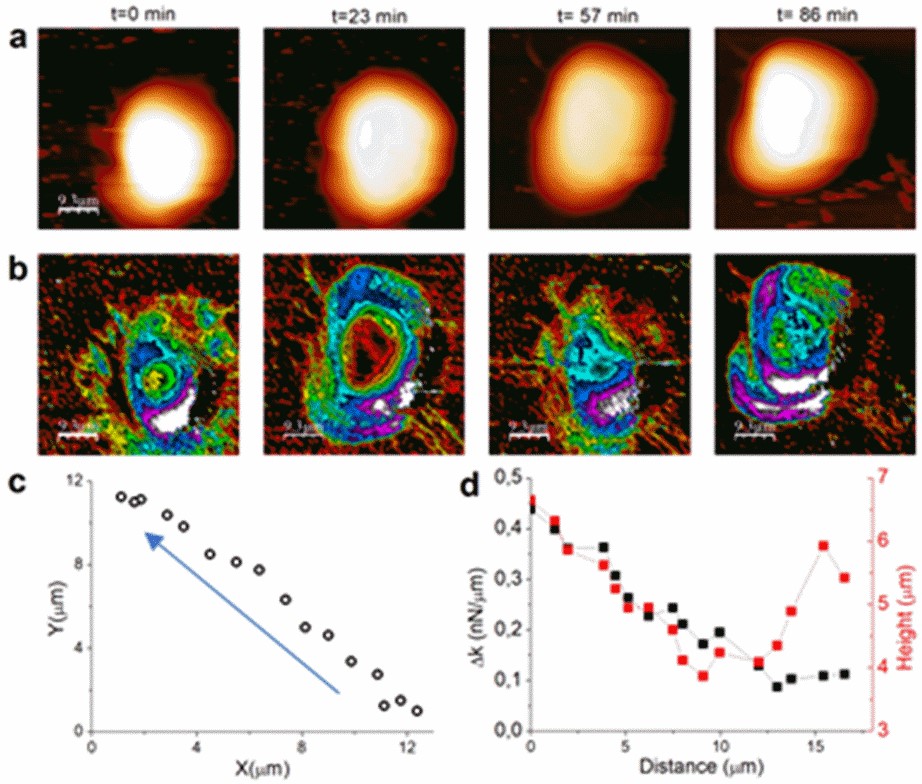

### Supplementary Figure 4

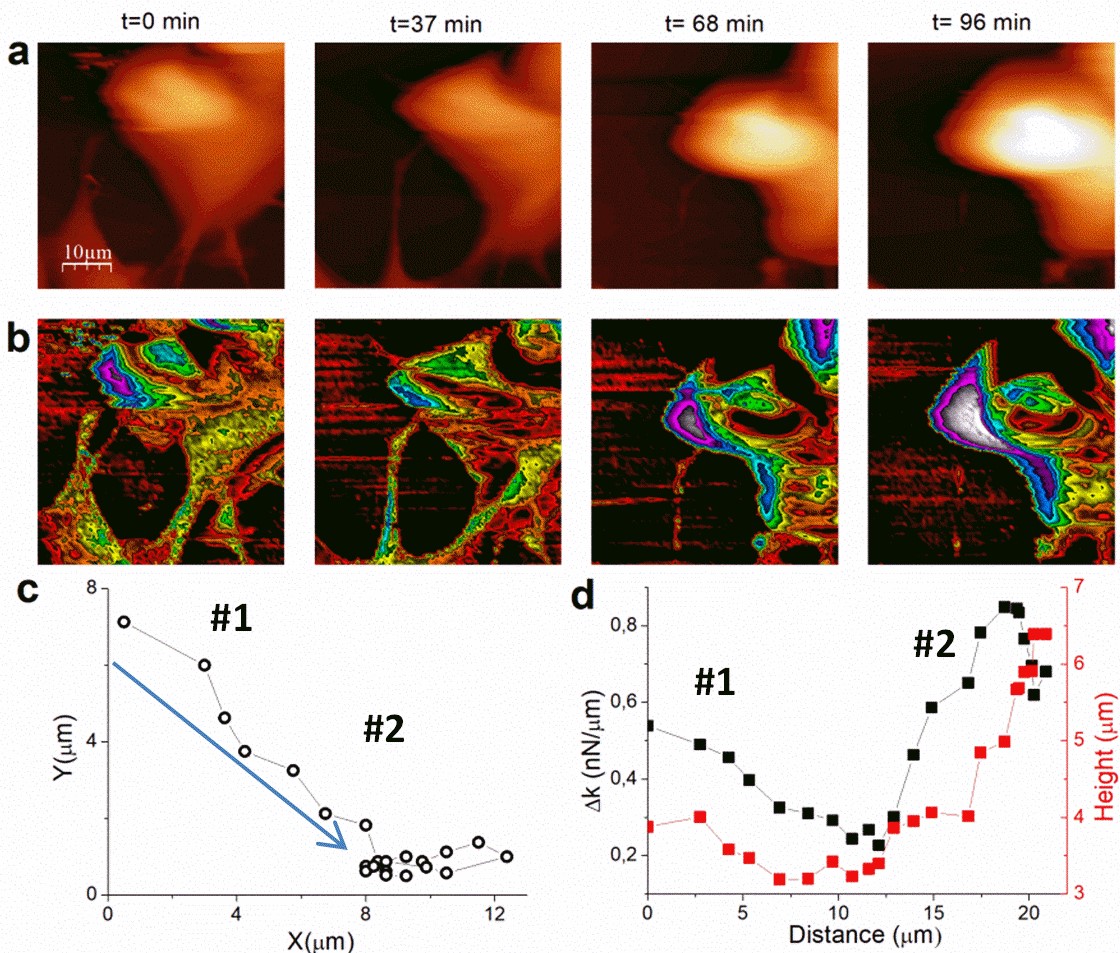

### Supplementary Figure 5

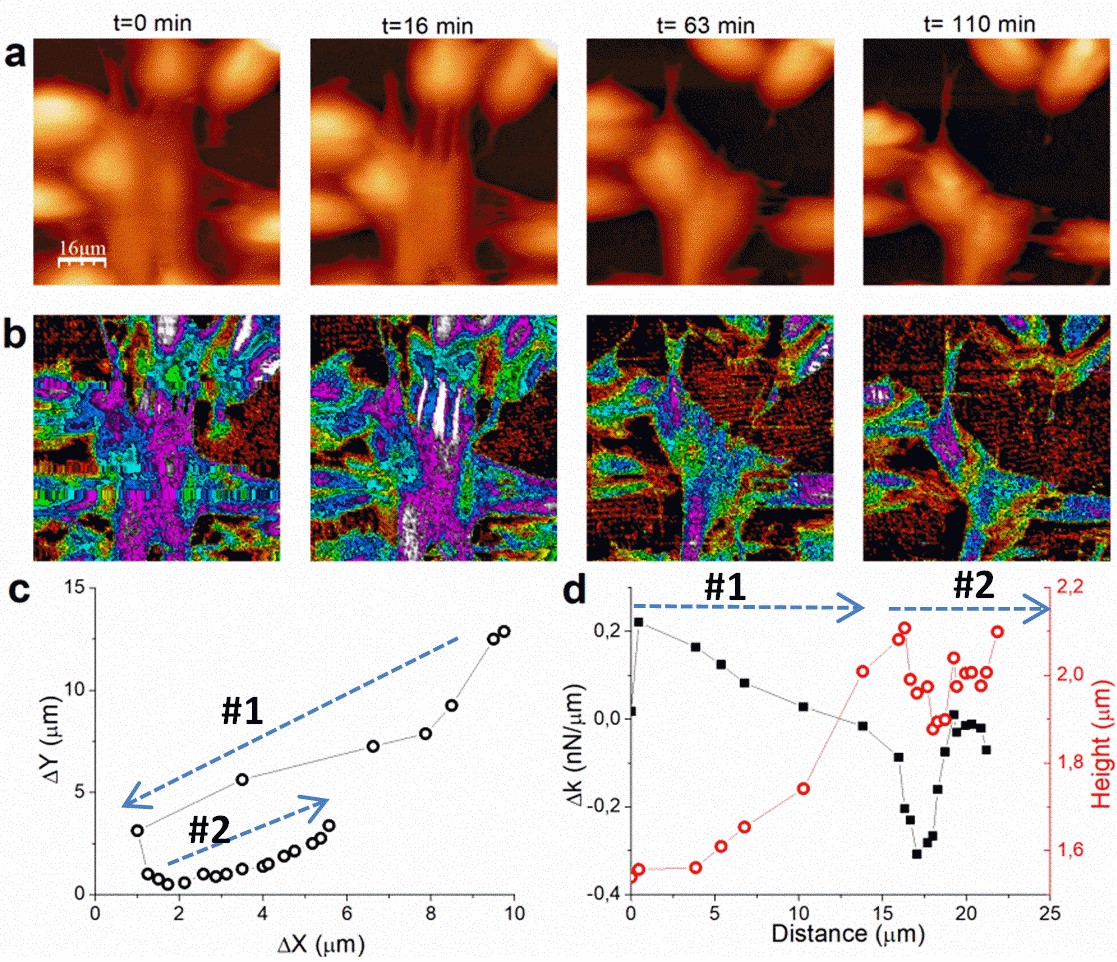
